# Glutamate and Dysconnection in the Salience Network: Neurochemical, Effective-connectivity, and Computational Evidence in Schizophrenia

**DOI:** 10.1101/828558

**Authors:** Roberto Limongi, Peter Jeon, Michael Mackinley, Tushar Das, Kara Dempster, Jean Théberge, Robert Bartha, Dickson Wong, Lena Palaniyappan

**Author notes:** Correspondence concerning this article should be addressed to Roberto Limongi, Robarts Research Institute, 1151 Richmond St. N, UWO, London, Ontario, Canada, N6A 5B7.

## Abstract

In the dysconnection hypothesis, psychosis is caused by NMDA hypofunction resulting in aberrant network connectivity. Combining a cognitive-control task, functional magnetic resonance spectroscopy, and functional magnetic resonance imaging, we tested this hypothesis in the salience network of 20 first-episode psychosis (FEP) and 20 healthy control (HC) subjects. Across groups, glutamate concentration in the dorsal anterior cingulate cortex (dACC) was associated with higher and lower inhibitory connectivity in the dACC and in the anterior insula (AI) respectively. Crucially, glutamate concentration correlated negatively with the inhibitory influence on the excitatory neuronal population in the dACC of FEP subjects. Furthermore, aberrant computational parameters of the cognitive-control task performance were associated with aberrant inhibitory connections. Finally, the strength of connections from the dACC to the AI correlated negatively with severity of social withdrawal. These findings support a link between glutamate-mediated cortical disinhibition, deficits in effective connectivity, and computational performance in psychosis.

## Introduction

The glutamate hypothesis (1–4) has been a central focus of interest in the study of the psychopathology of schizophrenia for more than 20 years. It states that dysfunction of glutamatergic neurotransmission is associated with signature negative symptoms of schizophrenia such as poor executive control and social withdrawal. The dysfunction of glutamatergic neurotransmission is likely caused by NMDA-receptor hypofunction (i.e., glutamate hypofunction) in inhibitory GABAergic interneurons which would lead to an increase in the synaptic gain of excitatory neurons (the disinhibition hypothesis, 5, 6). The abnormal increase of synaptic gain is also conceptualized as a disruption of the excitation-inhibition balance which, as described below, rests at the core of the dysconnection hypothesis (7) within the theoretical framework of “the Bayesian brain” (8).

The dysconnection hypothesis (c.f., disconnection hypothesis, 9)^1^ states that the psychopathology of schizophrenia should be studied at three levels of analysis: neurochemical, effective-connectivity (network-connectivity), and computational levels. At the computational level, the dysconnection hypothesis states that a *suboptimal* (e.g., schizophrenia) *Bayesian brain* (10) would overly afford confidence or precision (i.e., inverse variance) to its predictions about the external stimuli and would overestimate the reliability of the prediction errors (PE), leading to false inferences (e.g., hallucinations) and cognitive failures (e.g., cognitive control, 11). At the effective-connectivity level, a predictive coding algorithm (12), namely, hierarchical message passing between lower and higher cortical levels would be altered in terms of aberrant backward and forward interregional connectivity strength (13). Crucially, at the neurochemical level the aberrant connectivity strength would depend on increased synaptic gain in deep and superficial pyramidal cells, reflecting a decrease in the strength of intrinsic inhibitory connections, driven by NMDA hypofunction (7, 14). Therefore, the dysconnection hypothesis takes the glutamate hypothesis one step further by postulating that the disruption of excitation-inhibition balance makes itself evident at effective-connectivity level of analysis. In this work, we provide evidence in support of this hypothesis.

We provide evidence of the association between glutamate and dysconnection within the context of neurochemical, effective-connectivity, and computational levels of analysis. We will do this by studying the relationship between ^1^H-MRS glutamate in the dorsal anterior cingulate cortex ([Glu]_dACC_), the effective connectivity within the salience network, and cognitive-control dysfunction and negative symptoms in schizophrenia. Specifically, at the computational level we compare the performance of first-episode psychosis (FEP) and healthy control (HC) subjects in the Stroop task (15), which reliably engages the two nodes of the salience network (16, 17). Suboptimal Stroop computations in FEP are reflected in long reaction times and low response accuracy (18, 19). We show that these suboptimal computations are explained in terms of a drift-diffusion model as a specific case of a Bayesian decision making (20). At the effective-connectivity level, we show that the computational parameter associated with aberrant predictions in FEP maps onto forward and backward connections. Ultimately, we demonstrate at the neurochemical level that the dysconnection within the salience network is driven by [Glu]_dACC_. As we detail below, the salience network is an appropriate anatomical and functional motif (c.f., 21) to evaluate the relationship between all three levels of analysis.

The dACC is functionally specialized in conflict monitoring during the Stroop task (22). In the salience network, the right anterior cingulate cortex (dACC) is anatomically connected to the right anterior cingulate insula (AI) (23–25). Crucially, psychosis is associated with consistent structural deficits (26) as well as resting-state functional dysconnectivity in the salience network (27, 28). Given that the right AI is particularly sensitive to descending afferents from the right dACC (29), we expected the subtle variations of excitation-inhibition balance resulting from the putative glutamatergic abnormalities within the dACC to induce dysconnectivity in the salience network, affecting both intrinsic inhibitory connections of the dACC and the extrinsic connection with the AI. We anticipated the aberrant effective connectivity to account for aberrant prior beliefs and overly precise PE during cognitive conflict resolution and to the burden of negative symptoms of schizophrenia.

## Methods and Materials

### Participants

Twenty FEP subjects (8 female; age male M = 22.5, SD = 3.45; age female M = 22.37, SD = 3.07; age range [17 -28]) and 20 HC subjects (9 female, age male M = 21.64, SD = 4.27; female M = 21.67, SD = 3.04; age range [16 - 29]) participated in the study. No difference in age was detected across groups (p > |t| = 0.47). Subjects were recruited from the “Prevention and Early Intervention Program for Psychosis” in London, Ontario. Criteria for inclusion in the FEP group included (i) first clinical presentation with psychotic symptoms and (ii) Diagnostic and Statistical Manual of Mental Disorders (5th Edition) criteria A for schizophrenia satisfied. HC subjects did not report a personal history of mental illness or family history of psychotic disorders. The subjects’ consent was obtained according to the Declaration of Helsinki, and approval for the study was obtained from the University Human Ethics Committee for Health Sciences at the University of Western Ontario. Subjects were assessed for positive and negative symptoms of schizophrenia using the eight items of the Positive and Negative Syndrome Scale (PANSS-8, 30). All relevant demographic data are provided in the Supplemental Information. As detailed below, inside an MRI scanner, subjects underwent resting-state ^1^H-MRS, resting-state fMRI, and performed a color version of the Stroop task.

### ^1^H-MRS

We measured resting-state ^1^H-MRS during a four-minute block. The measurement was made in a Siemens MAGNETOM 7-Tesla MRI scanner using an 8-channel transmit/ 32-channel receive, head-only, radiofrequency coil at the Centre for Functional and Metabolic Mapping at the University of Western Ontario. A 2.0 x 2.0 x 2.0 cm (8cm^3^) ^1^H-MRS voxel was placed on the dACC where activation was expected based on our previous work (19) using the Stroop task (MNI: X = 1, Y = 16, Z = 38, Fig. 1). To locate the voxel, we used a two-dimensional anatomical imaging sequence in the sagittal direction (37 slices, TR = 8000 ms, TE = 70 ms, flip-angle (*α*) = 120°, thickness = 3.5 mm, field of view = 240×191 mm). Conventional ^1^H-MRS sequences use a short or “short-as-possible” echo time to minimize the T2 relaxation and J-coupling effects. However, a recent study by Wong, Schranz (31) proposed that a longer echo time for the semi-LASER sequence would improve glutamate measurement in the human brain. Therefore, one long echo time (100 ms) dataset was acquired using a semi-LASER ^1^H-MRS pulse sequence (repetition time = 7500 ms). A 32 channel-combined, water-suppressed spectral average was acquired using the semi-LASER ^1^H-MRS pulse sequence (repetition time = 7500 ms, echo time = 100 ms, N = 32). Water suppression was achieved using the VAPOR preparation sequence and a water-unsuppressed spectrum was also acquired for spectral post-processing. Spectral post-processing is described in the Supplemental Information.

**Fig. 1.**
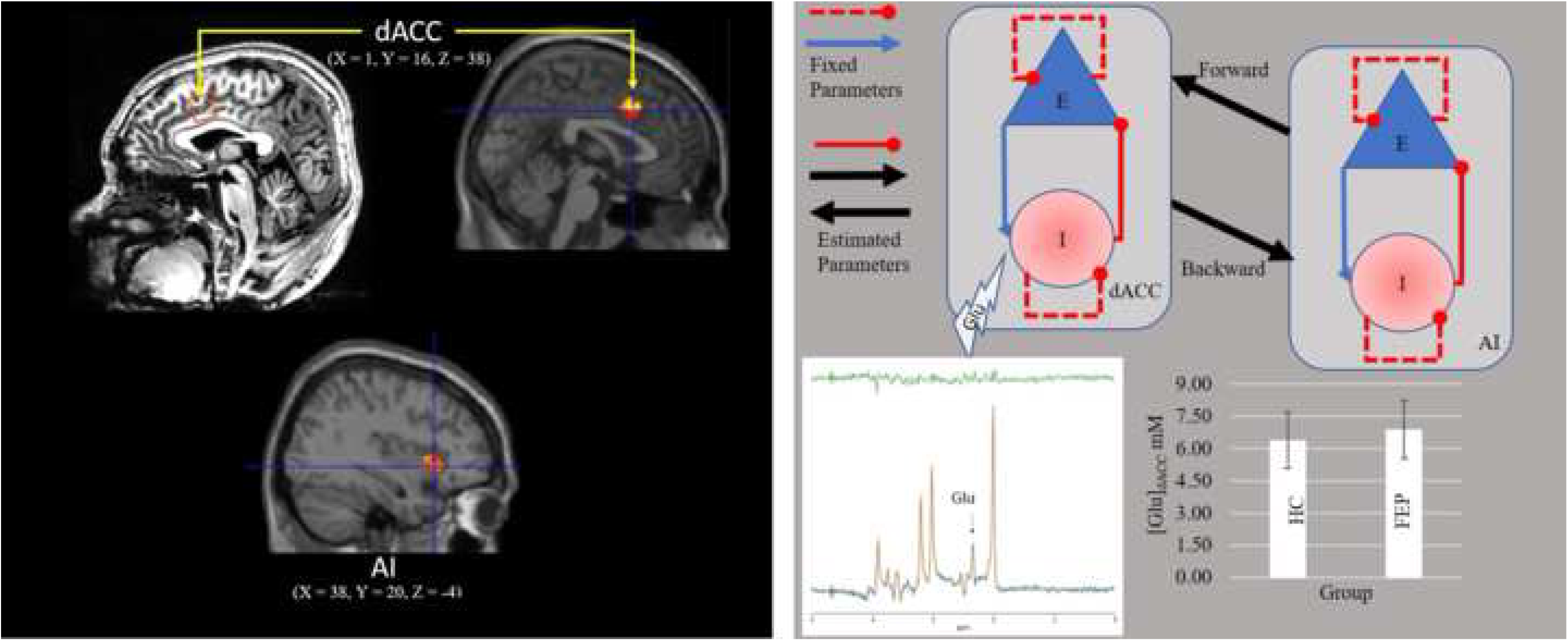
Voxel positioning for ^1^H-MRS measurement and two-neuronal-population DCM. Black box shows the voxel on the dACC for ^1^H-MRS (top) and the same voxel for fMRI (bottom). Grey box shows the two-neuronal-population DCM of the salience network. Each region comprises one population of excitatory neurons (E) and one population of inhibitory neurons (I). Parameters of effective connectivity represent the influence of inhibitory to excitatory connections (IE, assumed to be GABAergic neurons), the influence of excitatory to inhibitory connections (EI), the influence of self-inhibitory connections within each population (SE, SI), and the influence of excitatory population of one region on the excitatory population of the other region (EE, assumed to be glutamatergic connections). Whereas EI, SE, and SI parameters are fixed in the model, IE and EE are free parameters. Small white box shows a sample spectrum obtained using ^1^H-MRS semi-LASER (TE = 100 ms). Raw data are indicated in blue, fit spectrum is indicated in orange, and residual signal is shown above in green. Main glutamate signal contribution is indicated with an arrow (2.3 ppm). Since we aimed at demonstrating the relationship between the effect of [Glut]_dACC_ (observed measurements on the bar chart indicating no group difference) on IE and the ensuing consequences in the whole network, this two-state model was sufficient to test our hypothesis despite not capturing all the circuitry constraints of cortical columns (c.f., a four-neuronal-population model, 40).

### Resting-state fMRI

Using the same scanner as in the ^1^H-MRS measurement, we acquired 360 whole-brain functional images. A gradient echo-planar-imaging sequence was used with phase-encoding direction = A > > P, repetition time = 1000 ms, echo time = 20 ms, flip angle = 30 deg, field of view = 208 mm, field of view phase = 100 %, voxel dimension = 2 mm isotropic, slice thickness = 2 mm, multi-band acceleration factor = 3, acquisition time = 6 min. 26 s, and number of slices = 63 (interleaved slice order). Subjects were instructed to lie with their eyes open.

### Stroop task

We implemented the same Stroop task we used in Taylor, Neufeld (19). By pressing a key in a four-key response pad, subjects identified as quickly and as accurate as possible the color of the ink of a string of letters (on a gray background) representing either the name of a color (yellow, green, blue, or red) or a series of ‘Xs’ (‘XXXX’) projected on screen visible via a mirror located in the scanner’s coil. The task comprised four conditions: word-only (a word in white ink saying a color name, e.g., the word ‘yellow’ written in white ink), color-only (an ‘XXXX’ string with the color ink), congruent (the ink color matching the meaning of the word, e.g., the word ‘red’ in red ink), and incongruent (the ink color differing from the meaning of the word, e.g., the word ‘red’ written in blue ink). Subjects performed 20 trials per condition. They were allotted 2 s to respond with an interstimulus trial interval of 1 s during which a fixation cross was visible. Outside the scanner, each participant rehearsed the task until reaching 80 % accuracy — collapsed across all four conditions.

### Computational model of the Stroop performance

Based on our previous works (19), we were particularly interested in the incongruent condition in which we expected lower accuracy and longer reaction time in FEP than in HC. To verify this, we regressed accuracy and reaction time (separately) against condition, group, and the Condition × Group interaction via a mixed-effects linear model (32). Subjects were included as random effects.

We fit a hierarchical drift-diffusion model to the reaction time and accuracy data. In the hierarchical drift-diffusion model, subjects accumulate information and trigger a response after reaching an accumulation threshold. Formally, the hierarchical drift-diffusion model comprises four basic parameters representing the accumulation threshold, the starting point of the accumulation process, the accumulation (or drift) rate, and the non-decision processes such as stimulus encoding and motor execution (i.e., sensorimotor delay).

Predictive coding maps elegantly onto the hierarchical drift-diffusion model (11, 20, 33). Evidence accumulation corresponds to the accumulation of presynaptic afferent activity from neuronal populations encoding PE (i.e., superficial pyramidal cells), the drift rate represents the precision of the ascending PE, and the starting point represents the prior beliefs. Formally, corrected prior beliefs (i.e., a change in the starting point parameter) correlates with a steeper slope of the accumulation process (i.e., a larger absolute value of the drift rate indicating more precise PE).

Using Bayesian and Markov chain Monte Carlo methods (34), we estimated the parameters of the hierarchical drift diffusion model from subjects of each group separately. We assumed that subjects of a particular group were alike. Therefore, their parameters were constrained by their group parameters. This also allowed us to estimate between-groups difference without over or underestimating the true group-level variance (35). Prior distributions were informative as per the default option used in the HDDM package (34). The chain length was 200,000. The number of burn-in iterations was 2000, and the chains were generated with thinning = 20. We report the proportion of the estimate’s posterior distribution (PP) that differs from zero. To show differences between groups, we report the proportion of the posteriors in which the parameter estimate for one group differs from the other. Specifically, we expected larger drift rate (i.e., aberrant precision of PE) and larger starting point (aberrant prior beliefs) in the FEP group than in the HC. We did not have a-priori expectations about differences in the decision threshold and in non-decision processes. However, we report the differences in these parameters for completeness and post-hoc interpretation.

### Effective connectivity

#### Preprocessing and general linear model

Functional images were realigned, normalized to the Montreal Neurological Institute space, and spatially smoothed using a 4 mm (full width at half maximum) Gaussian kernel. We fit a general linear model to the images and included the six head movement parameters and the time series corresponding to the white matter and cerebrospinal fluid as regressors. In addition, we included a cosine basis set with frequencies ranging from 0.0078 to 0.1 Hz (36). Images were also high-pass filtered to remove slow frequency drifts (< 0.0078 Hz). By specifying an F-contrast across the basis set, we identified regions with blood oxygen level fluctuations within the range of frequencies specified above. On the *F*-contrast, a sphere (8-mm radius) was centered on the Montreal Neurological Institute coordinates corresponding to the dACC (the same coordinates that formed the centroid of the ^1^H-MRS voxel for the [Glu]_dACC_ measurements) and to the AI (X = 38, Y = 20, Z = -4, Fig. 1) —we selected these coordinates based on our previous results (37). At the peak voxel on each region of interest, we defined a sphere (8-mm radius) and extracted the timeseries (principal eigenvariate) summarizing the activity within the sphere.

#### Spectral dynamic causal model (DCM)

We estimated the resting-state effective connectivity within the salience network by fitting a two-state spectral DCM (38) to the resting-state fMRI data (39). Two-state DCM assumes two populations of neurons within a region: excitatory and inhibitory. For stability of the model, each sub-population comprises self-inhibition connections (which are fixed parameters). Crucially, two free parameters are fit to the fMRI data: interregional excitatory-to-excitatory connections and within-region inhibitory-to-excitatory connections (Fig. 1). Since we aimed at demonstrating the relationship between the effect of [Glu]_dACC_ on inhibitory-to-excitatory connections and the ensuing consequences in the entire network, this two-state model was sufficient to test our hypothesis despite not capturing all the circuitry constraints of cortical columns (c.f., a four-neuronal-population model, 40). For each participant, a fully connected DCM, with no exogenous inputs was specified and inverted using spectral DCM of SPM12 (https://www.fil.ion.ucl.ac.uk/spm/software/spm12/).

After inverting subjects’ DCMs, we estimated a series of parametric empirical Bayes models (41, 42) aiming to test, at a group level, our hypothesis against five alternative hypotheses. We fit six (general linear) models to the posterior estimates of effective connectivity from each subject. Our main hypothesis stated that [Glu]_dACC_ hypofunction would cause aberrant effective connectivity in the salience network which would account for aberrant prior beliefs and aberrant precision of PE. We tested this hypothesis by comparing the evidence supporting a “three-level” model with the evidence supporting five alternative models. The three-level (neurochemical, effective-connectivity, and computational) model comprised the following four covariates: (i) group, (ii) [Glu]_dACC_, (iii) precision of PE, and (iv) prior beliefs. Crucially, the model included the interactions between [Glu]_dACC_, prior beliefs, and precision of PE with Group. Each covariate was mean centered, and we used effect coding for groups (HC = -1, FEP = 1). All but the “behavioral model” (see below) were reduced versions of the three-level model.

A “no-glutamate” model represented the hypothesis that glutamate did not affect the salience network, the model did not include the main effect of [Glu]_dACC_ and its interaction with group. If [Glu]_dACC_ did not exert an effect on the effective connectivity of the salience network, the no-glutamate model would perform better than the three-level model. Similarly, if the computational parameters representing prior beliefs and precision of PE were not affected by the aberrant connectivity within the salience network, a model without these effects would perform better than the three-level model. We fit this alternative model and referred to it as the “no-computational” model. Alternatively, it could be that a model including the observed behavioral responses would perform better than a model comprising the computational parameters. We fit this model by substituting (in the three-level model) the behavioral responses and their interactions with group for the hierarchical drift-diffusion model parameters. We referred to this model as the “behavioral model”. For completeness, we fit a “group model” and a “null model”. The group model represented the hypothesis that changes in the salience network would be caused only by differences between groups. This model comprised only the main effect of group. The null model included only a single column of “ones” in the design matrix representing the hypothesis that neither groups nor the covariates were associated with the effective connectivity of the network.

To adjudicate between models, we performed Bayesian model comparison. At a group level, we first report the mean posterior estimate (collapsed across groups) of the strength of each connection along with the PP of models with each parameter, relative to models without. To evaluate our ‘three level’ hypothesis, we report the effect sizes; i.e., parameters β of the between-subject parametric empirical Bayesian model associated with each main effect and their interactions with group.

### Goodness of fit, convergence, and acquisition quality assessments

In the hierarchical drift-diffusion model, goodness of fit was assessed via posterior predictive checks (43). Convergence was assessed by computing the R-hat statistic (44). Proportion of variance accounted for by the dynamic causal models are reported in the Supplemental Information (Table S1). Finally, to assess the quality of our acquired ^1^H-MRS data, signal-to-noise ratio (SNRNAA) was calculated by dividing the amplitude of the NAA CH3 peak in the frequency domain by the standard deviation of the noise in the last 32 most upfield points of the spectrum. The linewidth was also calculated using water-unsuppressed spectra and measuring the FWHM of the water peak in Hertz (Hz).

**Table 1.** Summary statistics of accuracy and reaction time in the Stroop task

| Group | Condition | Accuracy (proportion) |  | Reaction Time (s) |  |
| --- | --- | --- | --- | --- | --- |
|  |  | M | SD | M | SD |
| FEP | Color-only | 0.99 | 0.09 | 1.06 | 0.27 |
|  | Congruent | 0.98 | 0.14 | 0.98 | 0.27 |
|  | Incongruent | 0.92 | 0.27 | 1.19 | 0.31 |
|  | Word-only | 0.97 | 0.17 | 0.95 | 0.26 |
| HC | Color-only | 1.00 | 0.05 | 0.89 | 0.20 |
|  | Congruent | 1.00 | 0.05 | 0.82 | 0.21 |
|  | Incongruent | 0.98 | 0.14 | 1.04 | 0.25 |
|  | Word-only | 1.00 | 0.00 | 0.79 | 0.18 |

### Data availability

Data supporting the findings of this study are available at https://uwoca-my.sharepoint.com/:f:/g/personal/rlimongi_uwo_ca/EmDE6zsduKpIinRzEhWoyz8B2OX83970WxL8KnG0dUj6IA?e=0ZxdO6.

## Results

### [Glu]_dACC_ does not differ across groups

Glutamate MR Spectroscopy assessment could not be performed in one FEP subject. Data from this subject were not analyzed. A frequentist two-sample t-test did not reveal statistically significant difference between the mean [Glu]_dACC_ in the FEP group (M = 6.90 mM, SD = 1.35) relative to the HC group (M = 6.38 mM, SD = 1.3); t_(36.72)_ = 1.21, p = 0.88 (Fig. 1).

### [Glu]_dACC_ does not account for between-groups difference in (Stroop) computational performance

Both FEP and HC subjects performed the task as instructed. Table 1 shows the summary statistics. The mixed-effects models confirmed the expected behavioral results (Table 2). Across conditions, FEP subjects performed worse than HC subjects. Planned t-tests confirmed that they were less accurate (t(_502_) = 3.71, p < 0.001, Cohen’s d = 0.28) and took longer (t(_674_) = 6.99, p < 0.001, Cohen’s d = 0.52) when resolving cognitive conflicts in the incongruent condition.

**Table 2.** Mixed-effect models of response accuracy and reaction time in the Stroop task

|  | Parameter | Estimate | SE | DF | t | P | Lower<br>95 % | Upper<br>95% |
| --- | --- | --- | --- | --- | --- | --- | --- | --- |
| Accuracy | Intercept | 0.979 | 0.006 | 36.69 | 156 | <.0001 | 0.966 | 0.992 |
|  | Group [FEP] | -0.015 | 0.006 | 36.69 | -2.32 | 0.0261 | -0.027 | -0.002 |
|  | Condition [Color-only] | 0.015 | 0.004 | 3000.87 | 3.71 | 0.0002 | 0.007 | 0.023 |
|  | Condition [Congruent] | 0.010 | 0.004 | 3000.86 | 2.33 | 0.0201 | 0.002 | 0.018 |
|  | Condition [Incongruent] | -0.030 | 0.004 | 3001.31 | -7.2 | <.0001 | -0.038 | -0.022 |
|  | Group [FEP] × Condition [Color-only] | 0.011 | 0.004 | 3000.87 | 2.79 | 0.0053 | 0.003 | 0.020 |
|  | Group [FEP]× Condition [Congruent] | 0.006 | 0.004 | 3000.86 | 1.4 | 0.1624 | -0.002 | 0.014 |
|  | Group [FEP] × Condition [Incongruent] | -0.016 | 0.004 | 3001.32 | -3.85 | 0.0001 | -0.024 | -0.008 |
| Reaction<br>Time | Intercept | 0.971 | 0.022 | 36.83 | 45 | <.0001 | 0.927 | 1.014 |
|  | Group [FEP] | 0.083 | 0.022 | 36.83 | 3.86 | 0.0004 | 0.040 | 0.127 |
|  | Condition [Color-only] | 0.011 | 0.007 | 2999.88 | 1.66 | 0.0968 | -0.002 | 0.024 |
|  | Condition [Congruent] | -0.069 | 0.007 | 2999.87 | -10.4 | <.0001 | -0.082 | -0.056 |
|  | Condition [Incongruent] | 0.155 | 0.007 | 2999.98 | 23 | <.0001 | 0.141 | 0.168 |
|  | Group [FEP] × Condition [Color-only] | 0.004 | 0.007 | 2999.88 | 0.53 | 0.5943 | -0.010 | 0.017 |
|  | Group [FEP]× Condition [Congruent] | 0.000 | 0.007 | 2999.87 | 0.06 | 0.9561 | -0.013 | 0.013 |
|  | Group [FEP] × Condition [Incongruent] | -0.004 | 0.007 | 2999.98 | -0.63 | 0.5271 | -0.017 | 0.009 |

The hierarchical drift-diffusion model showed that all parameter estimates in both groups differed from 0 (PP = 1.0). Subject-wise estimates are provided in the Supplemental Information (Table S2). Fig. 2 shows that the drift-rate parameter in the HC group (M = 2.83, SD = 0.18) was larger than in the FEP group (M = 1.83, SD = 0.2), proportion of the group’s posteriors in which the parameters differ (ΔPP) = 0.99. Furthermore, the starting point parameter in the FEP group was larger (i.e., closer to the decision boundary, M = 0.44, SD = 0.03) than in the HC group (M = 0.32, SD =0.03), ΔPP = 0.99. The decision threshold was lower in the FEP group (M = 2.06, SD = 0.32) than in the HC group (M = 2.52, SD = 0.09), ΔPP= 0.96. Finally, the non-decision processes took longer in the FEP group (M = 0.64, SD = 0.11) than in the HC group (M = 0.46, SD = 0.09), ΔPP = 0.99.

**Fig. 2.**
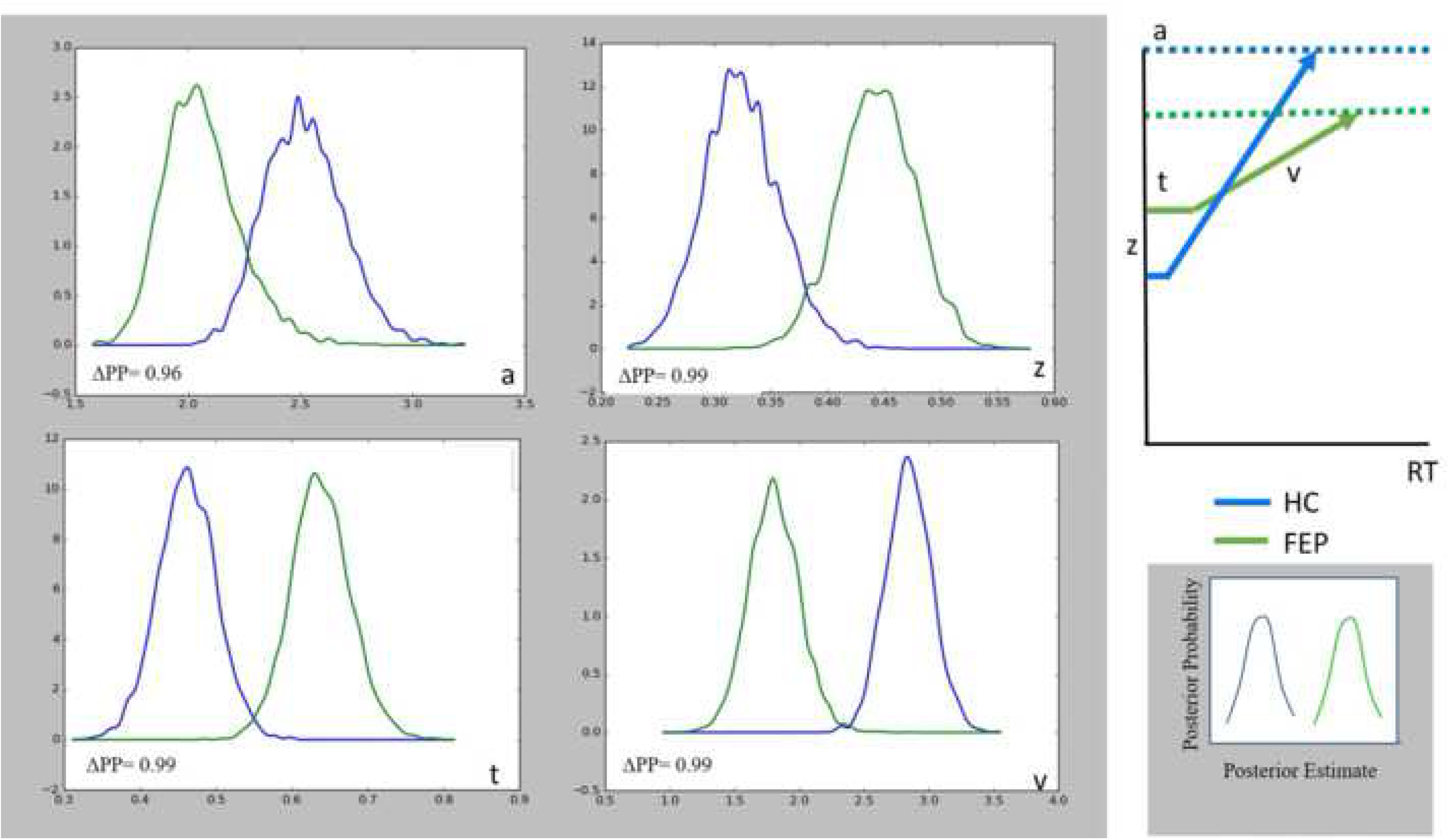
Parameter estimates of the hierarchical drift diffusion model. Top right, visual depiction of the parameter estimates of the drift diffusion model (z = starting point or prior beliefs, v = drift rate or precision of PE, t = non-decision processes, a = decision threshold, RT = reaction time). Left, proportion of the posteriors in which the parameter estimate for one group differs from the other (ΔPP).

To investigate whether [Glu]_dACC_ could explain between-groups differences in precision of PE and prior beliefs, we regressed the drift-diffusion model parameters on group, [Glu]_dACC_, and the [Glu]_dACC_ × Group interaction. Regarding prior beliefs, we found main effect of group, β = -0.06, SE = 0.0002, t(35) = -350, p < .0001, 95% CI [-0.059, -0.058] (congruent with the hierarchical drift-diffusion model results). However, we found no evidence for an effect of [Glu]_dACC_, β = 0.00006, SE = 0.0001, t(35) = 0.51, p = .61, 95% CI [-0.0002, 0.0003], nor [Glu]_dACC_ × Group interaction, β = -0.00009, SE = 0.0001, t(35) = -0.71, p = .48, 95% CI [-0.0003, 0.0002]. Similarly, regarding the precision of PE we found main effect of group, β = 0.5, SE = 0.05, t(35) = 9.11, p < .0001, 95% CI [0.39, 0.62] —also congruent with the hierarchical drift-diffusion model results— and we found no evidence for an effect of [Glu]_dACC_, β = -0.009, SE = 0.04, t(35) = -0.22, p = .83, 95% CI [-0.09, 0.08] nor [Glu]_dACC_ × Group interaction, β = 0.01, SE = 0.04, t(35) = 0.31, p = .76, 95% CI [-0.07, 0.098].

### [Glu]_dACC_ affects intrinsic inhibitory connections in the salience network

Bayesian model comparison revealed that the three-level model outperformed all the alternative models (Fig. 3). The mean estimates across groups revealed that in resting state, there was intrinsic inhibition in both regions (dACC, M = 2.25, PP =1.0; AI, M = 2.6, PP = 1.0) and an ensuing attenuation in the strength of excitatory connections between regions (dACC→AI, M = - 0.53, PP =1; AI→dACC, M = -0.43, PP =1.0).

**Fig. 3.**
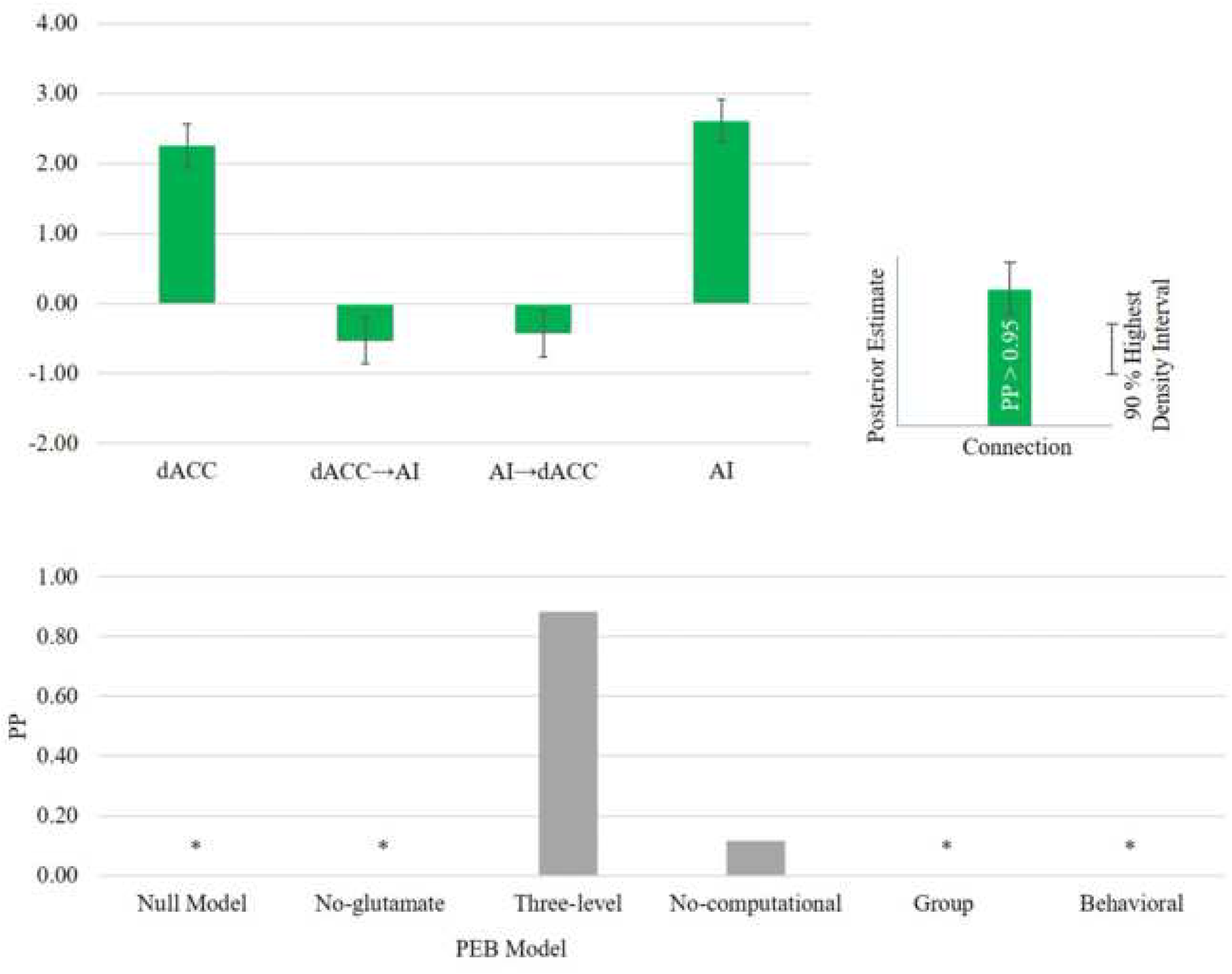
Bayesian model comparison results and mean parameter estimates in the salience network across groups. **\* PP ≈ 0.**

Fig. 4 shows no main effect of group on network parameters. However, main effect of [Glu]_dACC_ on self-connections in both the dACC and the AI was detected. The strength of intrinsic inhibition increased in the dACC (β = 0.09, PP = 0.98) and decreased in the AI (β = -0.07, PP = 0.96). Across groups, we did not find effect of prior beliefs on extrinsic connections. However, we found a main effect of the precision of PE on inhibitory connections. Across groups, the strength of inhibitory connections increased with the precision of PE (dACC, β = 0.9, PP = 1; AI, β = 0.85, PP = 1).

### [Glu]_dACC_ correlates negatively with intrinsic inhibitory influence in the dACC of FEP subjects, accounting for aberrant prior beliefs and aberrant precision of PE

Crucially, Fig. 5 shows that there was a Group × [Glu]_dACC_ interaction in inhibitory connections of the dACC. Connections were weaker in FEP than in HC (β = -0.11, PP =1.0). Furthermore, there was a Group × Prior-beliefs interaction. The effect of inhibitory connections on prior beliefs was stronger in FEP than in HC (dACC, β = 39.52, PP =1) (AI, β = 44.44, PP = 1). Surprisingly, the effect of excitatory connections on prior beliefs was weaker in FEP than in HC (dACC→AI, β = -9.13, PP =1; AI→dACC, β = -7.49, PP = 1). Finally, we found Group × PE precision interaction in inhibitory connections. The effect of these connections on the precision of PE was weaker in FEP (dACC, β = -1.56, PP = 1; AI, β = -1.08, PP = 1). In summary, [Glu]_dACC_ differentially affected the effective connectivity of the salience network, and the connectivity of the network was differentially associated with the parameter estimates of the hierarchical drift diffusion model in the presence of FEP.

**Fig. 4.**
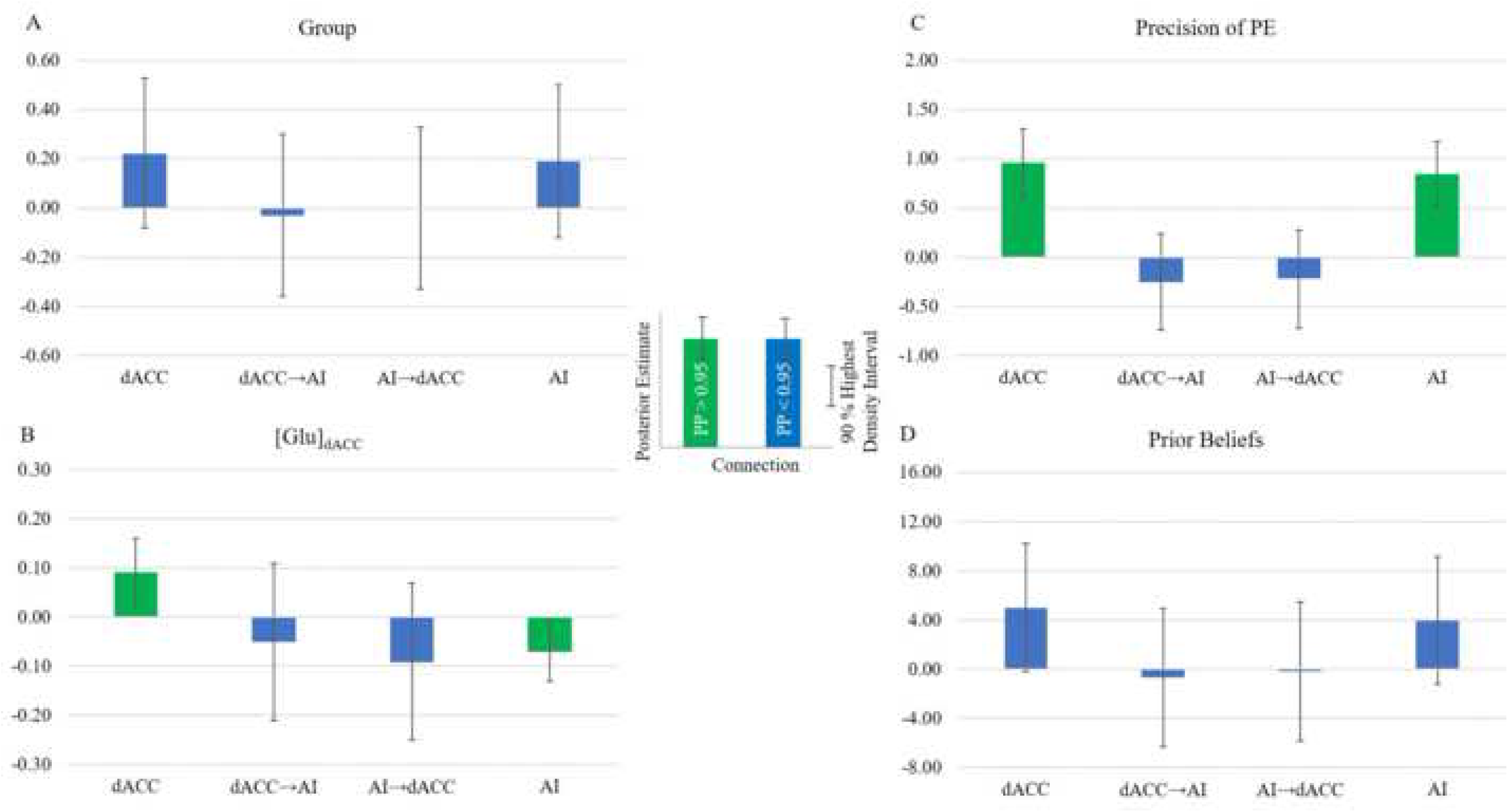
Main effects of the winning three-level parametric empirical Bayes model. In A, bars represent differences between groups as defined by the effect coding (FEP = 1, HC = -1). In B, C, and D, covariates were mean centered.

**Fig. 5.**
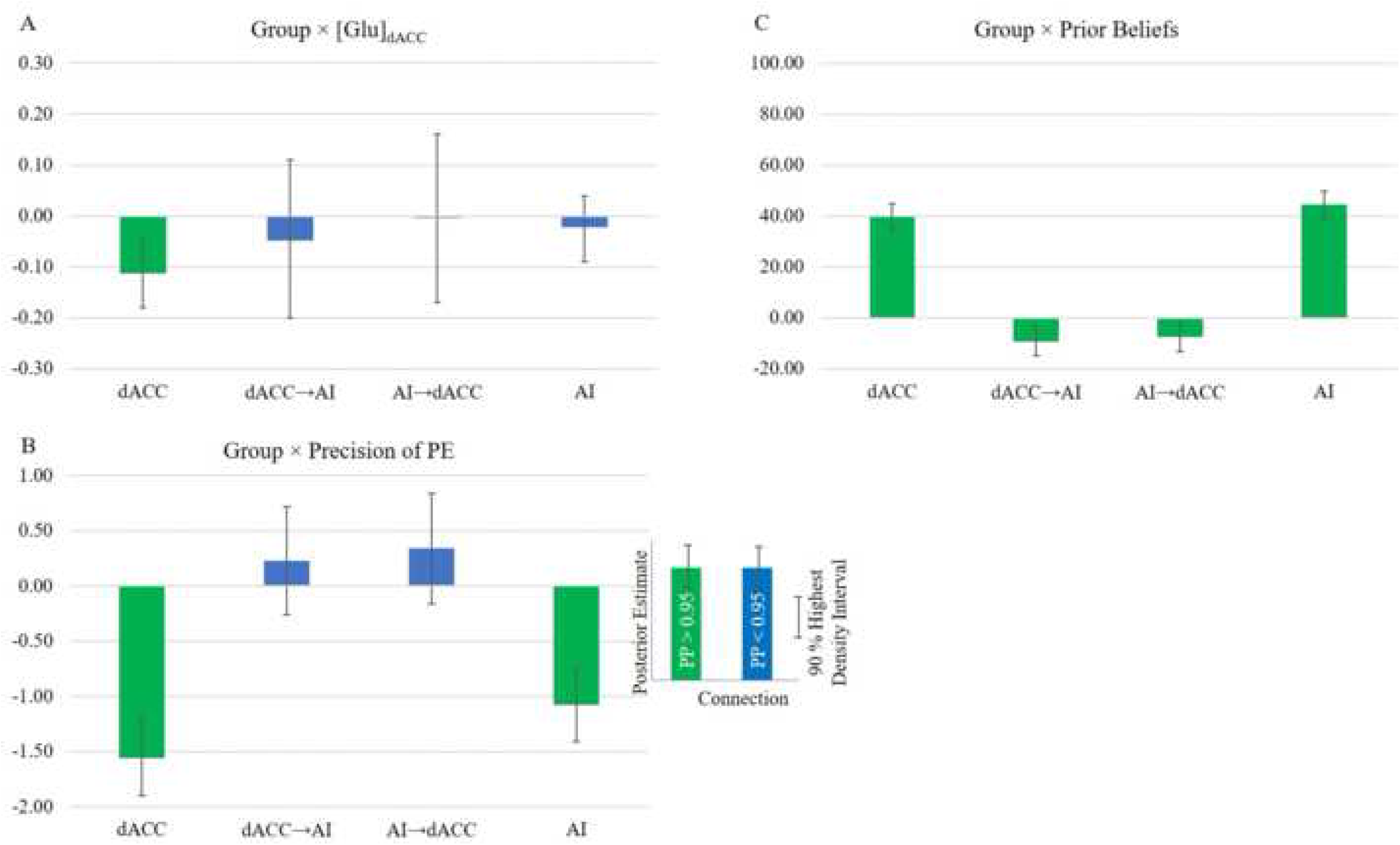
Interactions between group and covariates in the winning three-level parametric empirical Bayes model. In A, B, and C, positive values represent stronger effect in the FEP group, and vice versa.

### Connectivity strength correlates negatively with severity of social withdrawal

We assessed the association between the severity of negative symptoms (blunted affect, social withdrawal, and lack of spontaneity) and the strength of connections independently (i.e., three comparisons per connection) in the FEP group. Only the effect of backward connections on social withdrawal survived Bonferroni correction for the number of negative symptoms (three symptoms, P = 0.017). Severity of social withdrawal increased as the connectivity strength of dACC→AI connections decreased, β = -15.59, SE = 5.61, t(_16_) = 2.78, p = 0.013, 95% CI [27.43, 3.75] (Fig. 6).

**Fig. 6.**
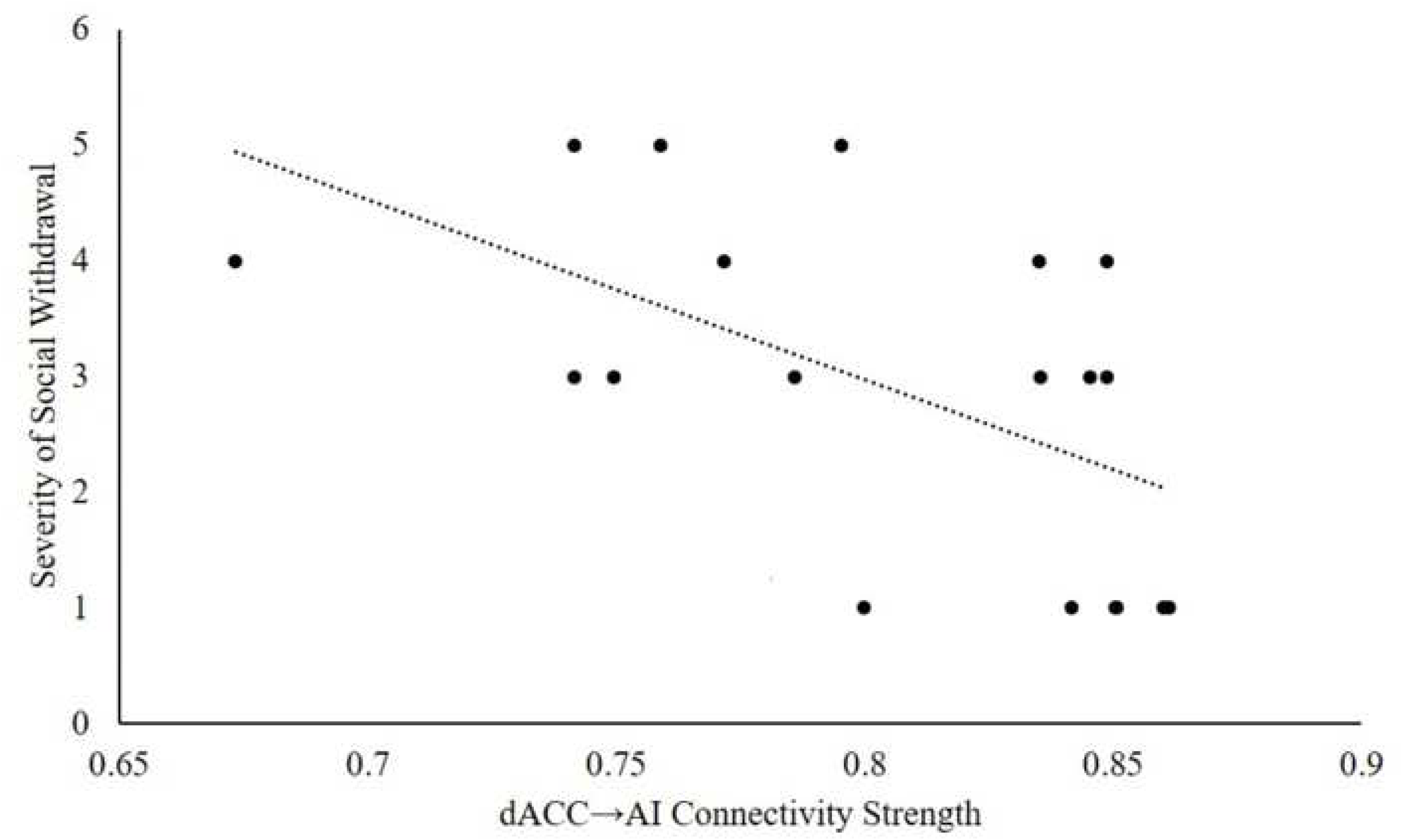
Association between Extrinsic Connectivity Strength and Severity of Social Withdrawal.

## Discussion

By using resting-state ^1^H-MRS, DCM of resting-state fMRI, and hierarchical drift-diffusion modeling, we have demonstrated that low computational performance during cognitive-conflict resolution in schizophrenia is explained by aberrant resting-state effective connectivity (dysconnection) in the salience network. This dysconnection was associated with glutamate hypofunction in the dACC, as proven by the fact that a model without the effect of [Glu]_dACC_ had low probability of having produced the fMRI data. [Glu]_dACC_ was associated with opposite intrinsic connectivity strength in both the dACC and the AI, indicating that [Glu]_dACC_ affected the entire network and confirming the sensitivity of the AI to subtle changes in the dACC excitation-inhibition balance. Crucially, in the FEP group the inhibitory influence on the excitatory population of the dACC decreased as a function of glutamate concentration. This finding is the first imaging evidence directly linking the glutamate hypofunction to the cortical disinhibition hypothesis in schizophrenia.

Using stochastic DCM, Bastos-Leite, Ridgway (45) found decreased intrinsic inhibition and decreased extrinsic connections in the default network of schizophrenia subjects. Similarly, using parametric empirical Bayes and DCM in a sample of schizophrenia subjects, Zhou, Zeidman (46) recently found decreased intrinsic inhibition and decreased extrinsic connectivity within the dorsal attention network. They speculated that both glutamate hypofunction and aberrant precision of prediction errors would be associated with this sort of dysconnectivity in large-scale networks. In the context of the salience network, we confirmed their hypothesis. Therefore, the fact that three independent groups have shown the same relationship between decreased intrinsic and extrinsic connections in the default, attention, and salience networks provide strong support to the dysconnection hypothesis of schizophrenia. Crucially, our results point to glutamate hypofunction and aberrant computations of sensory and prior precision as critical causes of dysconnection and are in line with recently reported independent preliminary data showing that the effect of glutamate hypofunction should be observed at the network level (47).

These results also demonstrate that establishing the relationship between glutamate hypofunction, the disinhibition hypothesis, and the pathophysiology of schizophrenia requires the integration of the neurochemical, effective-connectivity, and computational levels of analysis. This is supported by the fact that Bayesian model comparison afforded low probability to any model not comprising all three levels (PP ≈ 0.12, summed over all but the three-level model). Specifically, parameter estimates of resting-state aberrant connectivity (driven by resting-state [Glu]_dACC_) in the salience network accounted for computational parameters of cognitive dysfunction in schizophrenia. Furthermore, an increase in the effect of the inhibitory population on the excitatory population in both regions explains the precision subjects afforded to ascending information, in line with the known role of GABA interneurons on pyramidal circuits (48).

The three-level model also explains the compensatory effect of aberrant prior beliefs to aberrant precision of ascending information in FEP. At the computational level, FEP subjects needed to accumulate less information than HC (i.e., lower decision threshold), relied more on their prior beliefs (i.e., at the beginning of each trial they were closer to the decision boundary) than on sensory information, and were less cautious when resolving cognitive conflicts ―they tended to jump to conclusions (49). At a group level, we did not find evidence for a larger precision of PE in the FEP group than in the HC group. On the contrary, the HC group showed a steeper drift rate. However, within the FEP group, over affordance of precision to PE was associated with a decrease in the strength of inhibitory connections —in both regions (Fig. 5-B). Crucially, an increase in the inhibitory activity was associated with an increase in prior beliefs (Fig. 5-C), suggesting (at the effective-connectivity level) that more precise prior beliefs compensate for the aberrant precision of PE —as predicted by the dysconnection hypothesis (7).

A three-level model could provide alternative explanations for previously reported findings. Specifically, Taylor, Neufeld (19) found no relationship between [Glu]_dACC_ and deficits in Stroop performance of FEP subjects. One possible reason for this finding might be the lack of both a computational model (e.g., hierarchical drift-diffusion model) and a network model (e.g., DCM). This is supported by the fact that we found no direct association between [Glu]_dACC_ on either behavioral performance (i.e., reaction time and accuracy) or hierarchical drift-diffusion model parameters. However, we demonstrated that glutamate hypofunction affected the whole salience network and the aberrant effective connectivity of this network did affect the computational performance.

The advantage of a “three-level approach” is also supported by recent findings of Shaw, Knight (14). They used a four-neuronal-population DCM to model inhibitory (GABA) connections to superficial pyramidal cells. Their DCM parameters were associated with the behavioral performance. Our Bayesian model comparison showed that a model including behavioral measurements rather than the computational parameters underperformed the three-level model —which included the computational parameters. The findings of Shaw, Knight (14), however, are in concert with our idea that establishing a relationship between synaptic dysfunction and behavior (i.e., symptoms) requires modeling effective connectivity (at least at the regional scale) that would be disrupted by the synaptic dysfunction. More in general, along with this recent work, the current results suggest that symptoms of schizophrenia emerge from the interaction between the three levels of analysis.

Our results and the idea of interactive emergence allow us to provide an alternative interpretation to recent findings of Valton, Karvelis (50), who reported a computational model of Bayesian learning in schizophrenia and HC. Unlike our work, their study did not find differences in the precision of prior beliefs or in the precision of sensory evidence across groups. However, the fact that they did not include the effective-connectivity level of analysis in their experimental design left open the possibility of a Group × Precision interaction on the effective connectivity. Our results regarding [Glu]_dACC_ support this alternative explanation. We did not find main effect of group on [Glu]_dACC_ (c.f., no main effect of group on precision in 50). However, the effect of [Glu]_dACC_ on inhibitory connections in the dACC varied with group.

Valton, Karvelis (50) also proposed that prior beliefs in schizophrenia subjects might not influence behavioral performance in the absence of sensory stimulation. During resting state fMRI, our FEP subjects did not expect sensory stimulation. Interestingly, the weaker the connectivity strength in descending connections (representing prior beliefs) were in resting state the more biased were their responses towards the decision threshold. Heuristically, this could be interpreted as an increase in the confidence FEP subjects afforded to their prior beliefs during task performance, caused by a decreased in the strength of backward connections. Therefore, it is possible that the effect that prior beliefs might exert on behavioral performance might depend not only on the sensory stimulation (as suggested by 50) but also on the resting-state connectivity strength of descending connections which in hierarchical message passing represent prior beliefs.

It is intriguing that the relationship between the strength of extrinsic connections and the computational parameters of precision-weighted PE (and prior beliefs) does not conform to the assumption of an asymmetric flow of information under predictive coding. Canonical formulations of predictive coding would predict an increase in the strength of ascending connections associated with aberrant precision of PE and an increase in the strength of descending connections associated with compensatory aberrant prior beliefs. However, we found that the extrinsic connectivity strength did not reflect the precision of PE, whereas weak bidirectional connections correlated with prior beliefs. These data provide initial evidence of “bidirectional correlates of effective connectivity strength with prior beliefs” in hierarchical message passing with, as we elaborate upon below, implications for the underlying predictive coding theory of brain function and its translational corollary, the dysconnection hypothesis.

Initial works on predictive coding proposed an asymmetrical functional architecture for hierarchical message passing whereby prior beliefs would be conveyed by descending connections and PE would be conveyed by ascending connections. Not by coincidence, the connectivity asymmetry was motivated on the anatomical asymmetry widely described in early sensory areas (13). However, this scheme is not congruent with the effect of weak bidirectional connections on prior beliefs that our findings suggest.

A possible explanation of our findings is that the parameter estimates of the current DCM could be reflecting the activity of another type of excitatory inputs —for example, Von Economo neurons. DCMs do not differentiate this type of excitatory influence from others. This is a limitation of the current study that could be resolved in the future by incorporating additional excitatory neuronal populations and parameters of interlaminar connectivity (40).

We posit that the salience network might defy the canonical asymmetry of predictive coding due to the presence of Von Economo neurons, emerging as a network functionally specialized in the bidirectional processing of prior beliefs. Indeed, the “unique” role of Von Economo neurons in predictive coding has been recently suggested (51). However, asymmetric message passing does not contradict the principles of functional specialization and functional integration. For example, classic theories of working memory have been recently updated by integrating the principle of hierarchical message passing with the widely accepted functional specialization of the prefrontal cortex (52). Working memory, in the prefrontal cortex, has been recently regarded as a locus for the generation and maintenance of prior beliefs (53) which would be passed along to lower areas in the cortical hierarchy (54). Therefore, it is possible that the prefrontal cortex generates prior beliefs and deploys such beliefs downstream to the salience network. As a hub, the salience network would then functionally specialize in broadcasting prior beliefs (i.e., predictions) to lower sensory areas and interoceptive channels. The salience network would draw upon *bidirectional* circulation of prior beliefs to execute this functional integration.

The possibility of a functionally specialized two-node network for the processing of prior beliefs is further supported both by previous works on social isolation and suicidal behavior. For example, a post-mortem study of schizophrenia subjects that died by suicide showed a larger density of Von Economo neurons in the cingulate cortex of these subjects, compared with schizophrenia subjects who died due to other causes (55). Interestingly, social isolation (i.e., social withdrawal) is strongly related with both the (aberrant) salience network (55, 56) and suicidal behavior (57). These previous works are in consonance with the relationship between dACC→AI connectivity and severity of social withdrawal we found in this work.

In conclusion, this work provides evidence to the hypothesis that the glutamate hypofunction relies on the disinhibition hypothesis and manifests itself in the effective connectivity of the salience network. The three levels of analysis provide compelling explanation to deficits in cognitive control and negative symptoms in schizophrenia. Finally, a bidirectional functional architecture of prior beliefs of the salience network could be the key to understand both its coordinating role with the central executive network (58) and its cardinal role in psychosis (26).

## Supplementary Information

**Table S1.** Subject-level values of proportion of variance accounted for the three-level parametric-empirical-Bayes model

| Subject | HC | FEP |
| --- | --- | --- |
| 1 | 5.1 | 21.44 |
| 2 | 6.21 | 9.48 |
| 3 | 18.23 | 37.41 |
| 4 | 32.19 | 20.83 |
| 5 | 32.55 | 20.13 |
| 6 | 11.97 | 18.03 |
| 7 | 31.64 | 3.6 |
| 8 | 15.28 | 8.37 |
| 9 | 19.77 | 5.39 |
| 10 | 12.18 | 31.89 |
| 11 | 5.76 | 38.74 |
| 12 | 21.81 | 46 |
| 13 | 24.01 | 27.92 |
| 14 | 6.48 | 31.96 |
| 15 | 30.61 | 36.86 |
| 16 | 27.6 | 30.61 |
| 17 | 11.82 | 9.82 |
| 18 | 7.61 | 6.09 |
| 10 | 15.3 | 26.32 |
| 20 | 8.51 |  |

Note. The mean proportion of variance accounted for by the model was 0.17 (SD = 9.73, 95 % CI [12.67, 21.79]) in the HC group and 0.23 (SD = 12.93, 95% CI = [16.44, 28.91]) in the FEP group. —a proportion of a least 0.17 of variance accounted for by the model (averaged across subjects) is considered acceptable (1).

**Table S2.** Parameter estimates of the hierarchical drift-diffusion model

| Group |  | Parameter |  |  |  |  |  |  |  |
| --- | --- | --- | --- | --- | --- | --- | --- | --- | --- |
|  |  | a |  | v |  | t |  | z |  |
|  |  | mean | std | mean | std | mean | std | mean | std |
| HC | 1 | 2.535 | 0.259 | 2.872 | 0.263 | 0.455 | 0.057 | 0.326 | 0.034 |
|  | 2 | 2.558 | 0.275 | 2.884 | 0.272 | 0.540 | 0.064 | 0.326 | 0.034 |
|  | 3 | 2.650 | 0.299 | 2.735 | 0.253 | 0.524 | 0.065 | 0.325 | 0.034 |
|  | 4 | 2.538 | 0.236 | 2.845 | 0.260 | 0.300 | 0.038 | 0.326 | 0.034 |
|  | 5 | 2.535 | 0.241 | 2.881 | 0.267 | 0.363 | 0.044 | 0.326 | 0.034 |
|  | 6 | 2.678 | 0.295 | 2.561 | 0.285 | 0.563 | 0.058 | 0.324 | 0.034 |
|  | 7 | 2.373 | 0.248 | 2.841 | 0.263 | 0.377 | 0.050 | 0.327 | 0.034 |
|  | 8 | 2.592 | 0.266 | 2.863 | 0.270 | 0.447 | 0.052 | 0.326 | 0.034 |
|  | 9 | 2.541 | 0.258 | 2.871 | 0.265 | 0.454 | 0.055 | 0.326 | 0.034 |
|  | 10 | 2.439 | 0.241 | 2.795 | 0.254 | 0.455 | 0.054 | 0.326 | 0.034 |
|  | 11 | 2.475 | 0.235 | 2.554 | 0.291 | 0.548 | 0.064 | 0.325 | 0.034 |
|  | 12 | 2.462 | 0.237 | 2.902 | 0.268 | 0.360 | 0.048 | 0.327 | 0.035 |
|  | 13 | 2.431 | 0.263 | 3.051 | 0.331 | 0.458 | 0.059 | 0.327 | 0.035 |
|  | 14 | 2.344 | 0.282 | 2.926 | 0.288 | 0.503 | 0.067 | 0.327 | 0.035 |
|  | 15 | 2.510 | 0.236 | 2.912 | 0.275 | 0.324 | 0.041 | 0.326 | 0.034 |
|  | 16 | 2.656 | 0.300 | 2.701 | 0.256 | 0.486 | 0.062 | 0.325 | 0.034 |
|  | 17 | 2.509 | 0.245 | 2.767 | 0.262 | 0.590 | 0.064 | 0.325 | 0.034 |
|  | 18 | 2.507 | 0.272 | 3.011 | 0.319 | 0.582 | 0.068 | 0.326 | 0.035 |
|  | 19 | 2.505 | 0.244 | 2.891 | 0.266 | 0.389 | 0.050 | 0.326 | 0.034 |
|  | 20 | 2.474 | 0.245 | 2.804 | 0.262 | 0.528 | 0.059 | 0.325 | 0.034 |
| FEP | 1 | 2.365 | 0.486 | 2.305 | 0.403 | 0.821 | 0.082 | 0.443 | 0.035 |
|  | 2 | 2.619 | 0.548 | 1.954 | 0.377 | 0.739 | 0.090 | 0.442 | 0.035 |
|  | 3 | 1.371 | 0.173 | 1.322 | 0.366 | 0.533 | 0.029 | 0.441 | 0.035 |
|  | 4 | 1.916 | 0.368 | 2.524 | 0.434 | 0.743 | 0.063 | 0.444 | 0.035 |
|  | 5 | 1.797 | 0.243 | 1.633 | 0.335 | 0.642 | 0.056 | 0.443 | 0.035 |
|  | 6 | 1.943 | 0.263 | 1.784 | 0.321 | 0.543 | 0.050 | 0.444 | 0.035 |
|  | 7 | 1.854 | 0.275 | 1.202 | 0.350 | 0.800 | 0.095 | 0.443 | 0.035 |
|  | 8 | 2.173 | 0.390 | 2.422 | 0.421 | 0.652 | 0.056 | 0.443 | 0.035 |
|  | 9 | 1.707 | 0.268 | 2.259 | 0.404 | 0.640 | 0.046 | 0.443 | 0.035 |
|  | 10 | 2.006 | 0.221 | 0.746 | 0.280 | 0.681 | 0.065 | 0.442 | 0.035 |
|  | 11 | 2.580 | 0.457 | 1.797 | 0.338 | 0.573 | 0.067 | 0.442 | 0.035 |
|  | 12 | 2.439 | 0.393 | 1.877 | 0.322 | 0.463 | 0.051 | 0.443 | 0.035 |
|  | 13 | 1.950 | 0.257 | 1.654 | 0.323 | 0.707 | 0.057 | 0.442 | 0.035 |
|  | 14 | 2.460 | 0.452 | 2.248 | 0.404 | 0.567 | 0.054 | 0.442 | 0.035 |
|  | 15 | 2.411 | 0.350 | 1.621 | 0.305 | 0.546 | 0.053 | 0.441 | 0.035 |
|  | 16 | 1.974 | 0.285 | 2.052 | 0.343 | 0.395 | 0.042 | 0.445 | 0.035 |
|  | 17 | 1.666 | 0.214 | 1.290 | 0.302 | 0.739 | 0.063 | 0.445 | 0.035 |
|  | 18 | 1.905 | 0.274 | 2.021 | 0.355 | 0.599 | 0.048 | 0.443 | 0.035 |
|  | 19 | 2.147 | 0.317 | 1.839 | 0.332 | 0.733 | 0.065 | 0.442 | 0.035 |

### Spectral post-processing

The spectral acquisition was analyzed to measure glutamate concentration using methods previously developed in our laboratory (2–5). Specifically, the 32 spectral acquisition was corrected for frequency and phase drifts as described in Near, Edden (6). The corrected spectra were then averaged into one spectrum to represent the entire four-minute resting-state block. The averaged spectrum underwent post-processing using combined QUALITY Eddy Current Correction (7), using 400 QUALITY (quantification improvement by converting lineshapes to the Lorentzian type) points, to reduce linewidth distortions as well as Hankel Singular Value Decomposition (HSVD) water removal to remove any residual water signal between 4.2 ppm and 5.7 ppm before being fit with fitMAN (2, 3), a time-domain fitting algorithm that uses a non-linear, iterative Levenberg-Marquardt minimization algorithm to echo time-specific prior knowledge templates. The metabolite fitting template included eighteen brain metabolites: alanine, aspartate, choline, creatine, γ-aminobutyric acid (GABA), glucose, glutamate, glutamine, glutathione, glycine, lactate, myo-inositol, Nacetyl aspartate, N-acetyl aspartyl glutamate, phosphorylethanolamine, scyllo-inositol, and taurine. As well, a single peak was used to fit the water unsuppressed spectrum.

Using Barstool (2), tissue-specific (gray matter, white matter, and CSF) T1 and T2 relaxations were corrected through partial volume segmentation calculations of voxels mapped onto T1-weighted images acquired using a 0.75 mm isotropic MP2RAGE sequence (repetition time = 6000 ms, TI1= 800 ms, TI2 = 2700 ms, flip-angle 1 (α1) = 4°, flip-angle 2 (α2) = 5°, field of view = 350 mm × 263 mm × 350 mm, acquisition time = 9 min. 38 s, iPATPE = 3 and 6/8 k-space). Finally, glutamate concentration quantification was calculated using the postprocessed water suppressed and unsuppressed spectra for each participant along with voxel-appropriate, tissue-specific relaxation time adjustments (8).

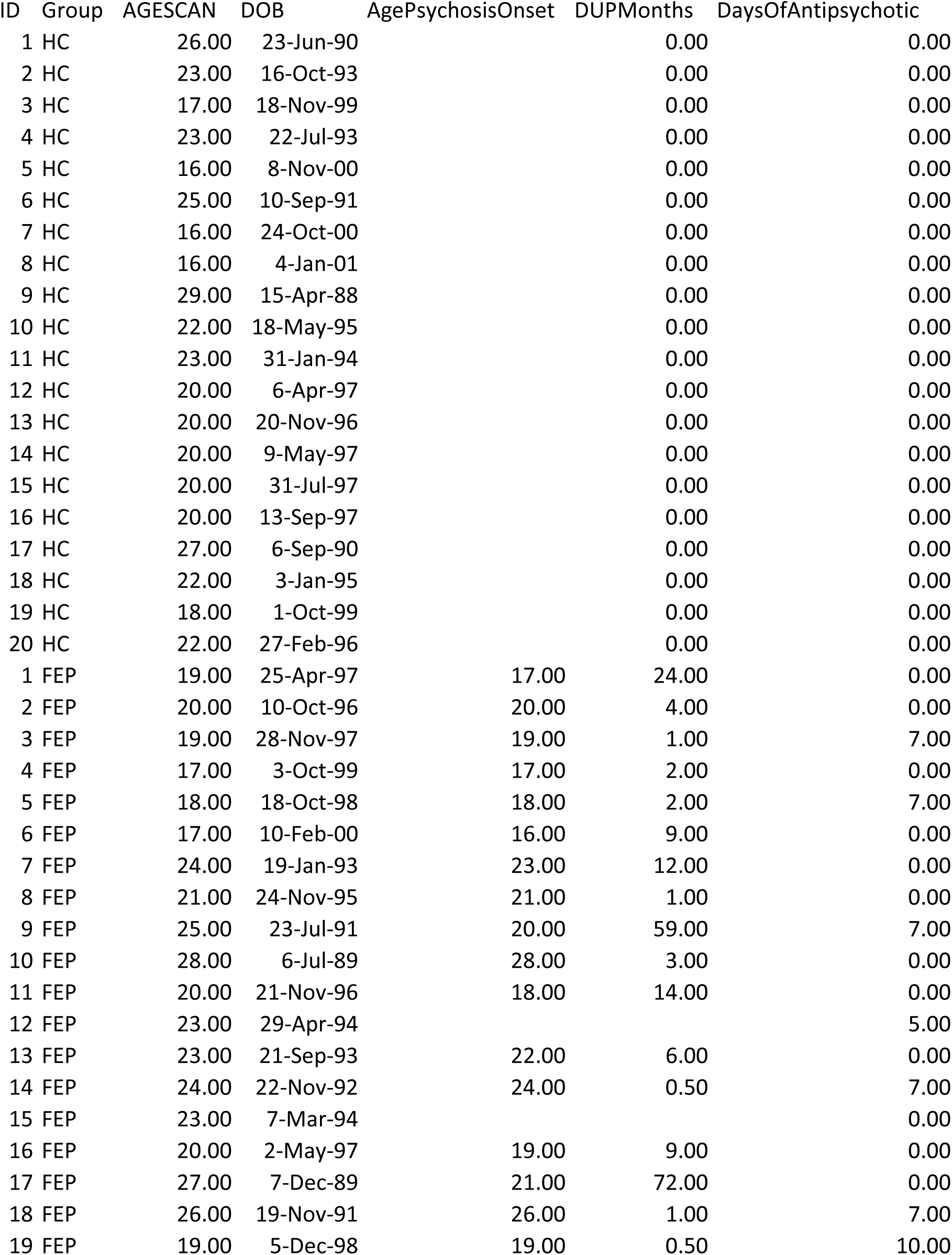
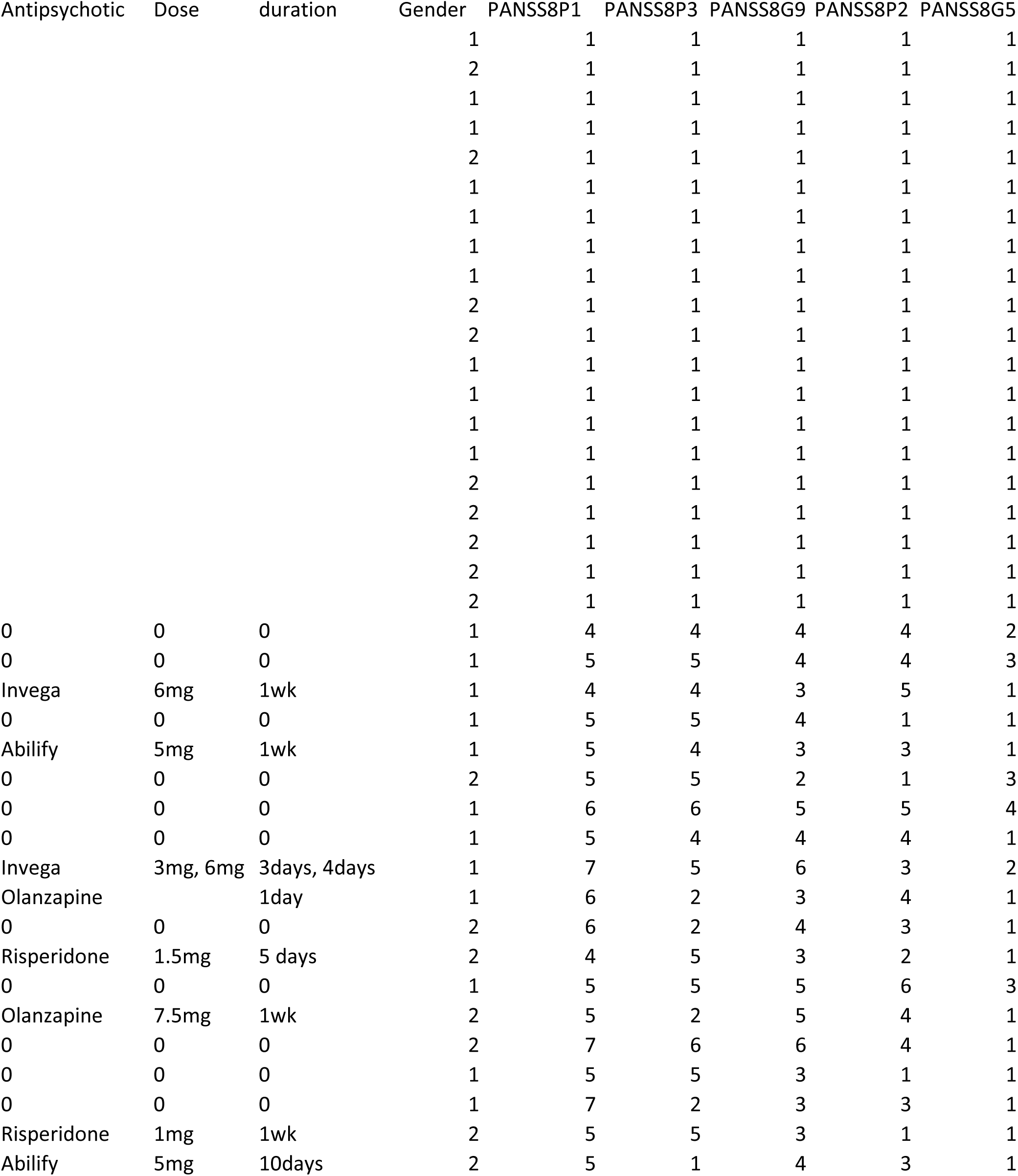
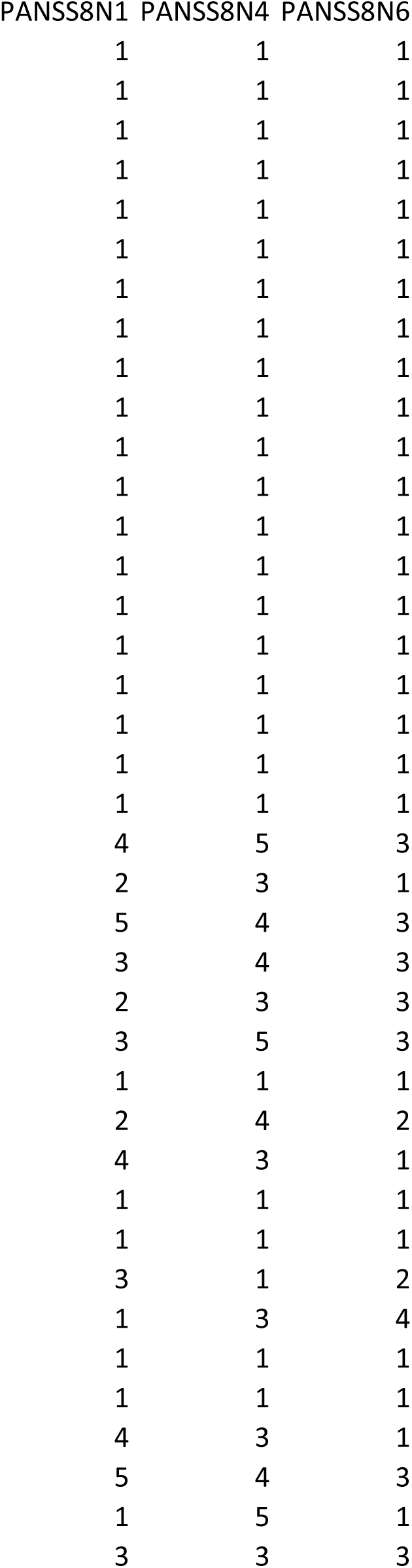

## Footnotes

1 Here, “dysconnection” means abnormal functional integration or effective connectivity. This meaning differs from the meaning of “disconnection” which refers to collapsed or disintegrated cognitive functions.

